# Nanosecond pulsed electric field precisely manipulates cytoplasmic Ca^2+^ oscillations during oocyte activation

**DOI:** 10.1101/2025.04.03.646992

**Authors:** Yi-Dan Sun, Tong An, Rong Liang, Yu-Wen Luo, Hong-Ze Xia, Lei Fu, Shuo Han, Yi-Xiao Zhu, Zi-Yi Song, Xue-Yan Bai, Yao Fu, Xiang-Wei Fu, Yun-Peng Hou, Qun Lu

**Affiliations:** State Key Laboratory of Animal Biotech Breeding, College of Biological Sciences, China Agricultural University; Beijing, China; Medical Center for Human Reproduction, Beijing Chao-Yang Hospital, Capital Medical University; Beijing, China; Department of Obstetrics and Gynecology, Peking University People’s Hospital; Beijing, China; National Engineering Laboratory for Animal Breeding, College of Animal Science and Technology, China Agricultural University; Beijing, China

**Keywords:** artificial oocyte activation, nanosecond pulsed electric field, Ca^2+^ oscillations, infertility treatment

## Abstract

Oocyte activation deficiency (OAD) is a primary cause of fertilization failure following intracytoplasmic sperm injection (ICSI), a problem that can potentially be overcome through artificial oocyte activation (AOA). However, concerns persist regarding the safety and efficacy of AOA techniques in clinical practice. In this study, nanosecond pulsed electric field (nsPEF) is proposed as a safe and controllable method for oocyte activation *in vitro* by precisely manipulating Ca^2+^ signaling in the ooplasm. Mouse oocytes collected from oviducts were exposed to nsPEF stimulation at varying intensities to induce distinct Ca^2+^ signaling patterns. Subsequently, these oocytes underwent parthenogenetic activation and were cultured to assess developmental potential up to the blastocyst stage. The sperm-initiated physiological Ca^2+^ oscillations are successfully mimicked by one series of Ca^2+^ signals induced by nsPEF at low or medium intensities, and improve activation efficiency and developmental potential compared to calcium ionophore A23187. Low-intensity nsPEF pulses mediated repetitive extracellular Ca^2+^ influx in a non-invasive, electro-permeable manner. Medium-intensity nsPEF stimulation triggered periodic Ca^2+^ release from the endoplasmic reticulum (ER) via the PIP_2_-IP_3_-IP_3_R pathway, producing physiological-like Ca^2+^ oscillations. The non-invasive nsPEF method ensures safe oocyte activation by preserving cellular integrity and minimizing stress responses. The efficacy of nsPEF in precisely manipulating Ca^2+^ signaling patterns was further validated in human unfertilized oocytes. Our findings present a novel AOA approach with enhanced safety and efficacy, especially for patients experiencing repeated fertilization failures. It is anticipated that nsPEF is emerging as a promising technology for addressing male infertility issues, and may offer a potential alternative fertility treatment, bypassing sperm for initiation of oocyte activation.

## 1. Introduction

The World Health Organization reports that approximately 17.5% of couples worldwide are affected by infertility.^1^ In vitro fertilization and embryo transfer (IVF-ET) remains a cornerstone of assisted reproductive technology (ART), with intracytoplasmic sperm injection (ICSI) serving as a specialized technique for severe male factor infertility or prior IVF failure.^2^ However, after implementing ICSI, some couples remain to face fertilization rates below 30%, and up to 3-5% of ICSI cycles may result in total fertilization failure (TFF), with a concerning recurrence in subsequent attempts.^3^ These repeated TFF after ICSI imposes significant physical and psychological distress on infertile couples.

In these cases of repeated TFF after ICSI, the majority are due to defective sperms, such as Globozoospermia, Teratozoospermia, Severe oligozoospermia, Non-obstructive azoospermia, etc^4^, which lead to varying degrees of oocyte activation deficiency (OAD).^5,6^ Sperm from such patients was linked with lacking or mutant phospholipase C zeta (PLCζ), the chief physiological inducer of intracellular calcium ions ([Ca^2+^]_i_) oscillations at fertilization. PLCζ promotes intracellular phosphatidylinositol 4,5-bisphosphate (PIP_2_) hydrolyze into inositol 1,4,5 triphosphate (IP_3_) to mediate the release of Ca^2+^ from the endoplasmic reticulum (ER).^7^ Then, the mechanisms that regulate intracellular calcium homeostasis, calcium-induced calcium release (CICR) and store-operated Ca^2+^ entry (SOCE), are activated by increased [Ca^2+^]_i_ level and induce periodic [Ca^2+^]_i_ oscillations in oocytes. The characteristic pattern of Ca^2+^ oscillations is crucial for oocyte activation and embryo development.^8,9^

Artificial oocyte activation (AOA) has been put forward to rescue oocytes that fail to be activated by defective sperms. This technique attempts to artificially induce the Ca^2+^ concentration increase in oocytes, which then assists the defective sperms to activate oocytes.^10^ Various oocyte activation methods have been applied in human ART, including chemical, mechanical, and physical methods.^3^ Among them, the chemical activation method, such as calcium ionophore (A23187), is commonly used in clinical practice.^11–13^ The activation rate, embryo quality, clinical pregnancy rate, and live birth rate of ICSI-AOA cycles are improved compared with the previous conventional ICSI cycle.^14–16^ Nevertheless, these activation methods can only induce a single or several Ca^2+^ waves rather than physiological sustained Ca^2+^ oscillation induced by sperm.^10,17^ The activation efficiency of ICSI-AOA varies ranging from 25% to 82.5% in different studies, and the embryo quality also varies greatly.^8,18,19^ Therefore, it seems to be urgent to seek AOA methods that quantitatively regulate [Ca^2+^]_i_ oscillations in oocytes, and explore how to induce physiological Ca^2+^ oscillations and their related biological mechanisms.

Electrical activation is a pretty important physical AOA method. The plasma membrane forms pores under the conventional electric field with durations from milliseconds to microseconds to induce extracellular Ca^2+^ flow into the cytoplasm.^20,21^ The human unfertilized oocytes completed the second meiosis and underwent early embryonic development after exposure to electrical stimulation.^22^ However, with the longer duration, the sizes of pores on the cell membrane are larger, which is difficult to recover and may even be irreversible, eventually lead to cell death ^23–25^, thus, the application of electrical activation in AOA treatment is limited.

Nanosecond pulsed electric field (nsPEF) with high-voltage directly impact structures of both plasma membrane and organelle membranes ^26–28^ to increase [Ca^2+^]_i_ concentration.^29^ Oocytes store substantial amounts of Ca^2+^ in organelles such as the ERs and mitochondria, which could undergo similar structural alterations in the biomembrane under nsPEF exposure.^30,31^ While this is clearly different from conventional electrical activation that only generates micropores in the plasma membrane. Considering the significance of Ca^2+^ oscillations in oocyte activation and embryonic development, greater attention should be directed towards how nsPEF stimulation regulate [Ca^2+^]_i_ change and formation of [Ca^2+^]_i_ oscillations akin to physiological conditions. However, the mechanism by which nsPEF influences intracellular and extracellular Ca^2+^ dynamics to activate oocyte development remains unclear.

This study aims to determine the optimal and safe parameter of nsPEF exposure for oocyte activation and embryo development. We also investigate the intrinsic mechanisms for the generation and maintenance of sustained Ca^2+^ oscillations. Through characteristic of nsPEF-induced [Ca^2+^]_i_ dynamics in human oocytes, we elucidate the nsPEF’s biophysical thresholds that ensure effective activation without compromising cytoplasmic integrity.

## 2. Results

### 2.1. Cytoplasmic Ca^2+^ signal patterns are quantitatively manipulated by nsPEF stimulation

The patterns of [Ca^2+^]_i_ signaling in response to nsPEF with different electric field intensities were represented in **Figure 1A-C**. After nsPEF stimulation at low intensity, oocytes exhibited a single and rapid [Ca^2+^]_i_ spike (Figure 1A). When the pulses intensity was raised to a moderate level, a specific sustained [Ca^2+^]_i_ oscillation was observed in the ooplasm, consisting of 5-8 Ca^2+^ spikes (Figure 1B), which had nearly the same shape as the spontaneous [Ca^2+^]_i_ oscillations at fertilization. At a relatively high field intensity, the pulse-induced Ca^2+^ wave displayed a swift [Ca^2+^]_i_ increase, followed by a slow decrease and gradual return to stable [Ca^2+^]_i_ concentration (Figure 1C), which was similar to the [Ca^2+^]_i_ wave induced by chemical activator A23187, as shown in Figure 1D.

**Figure 1.**
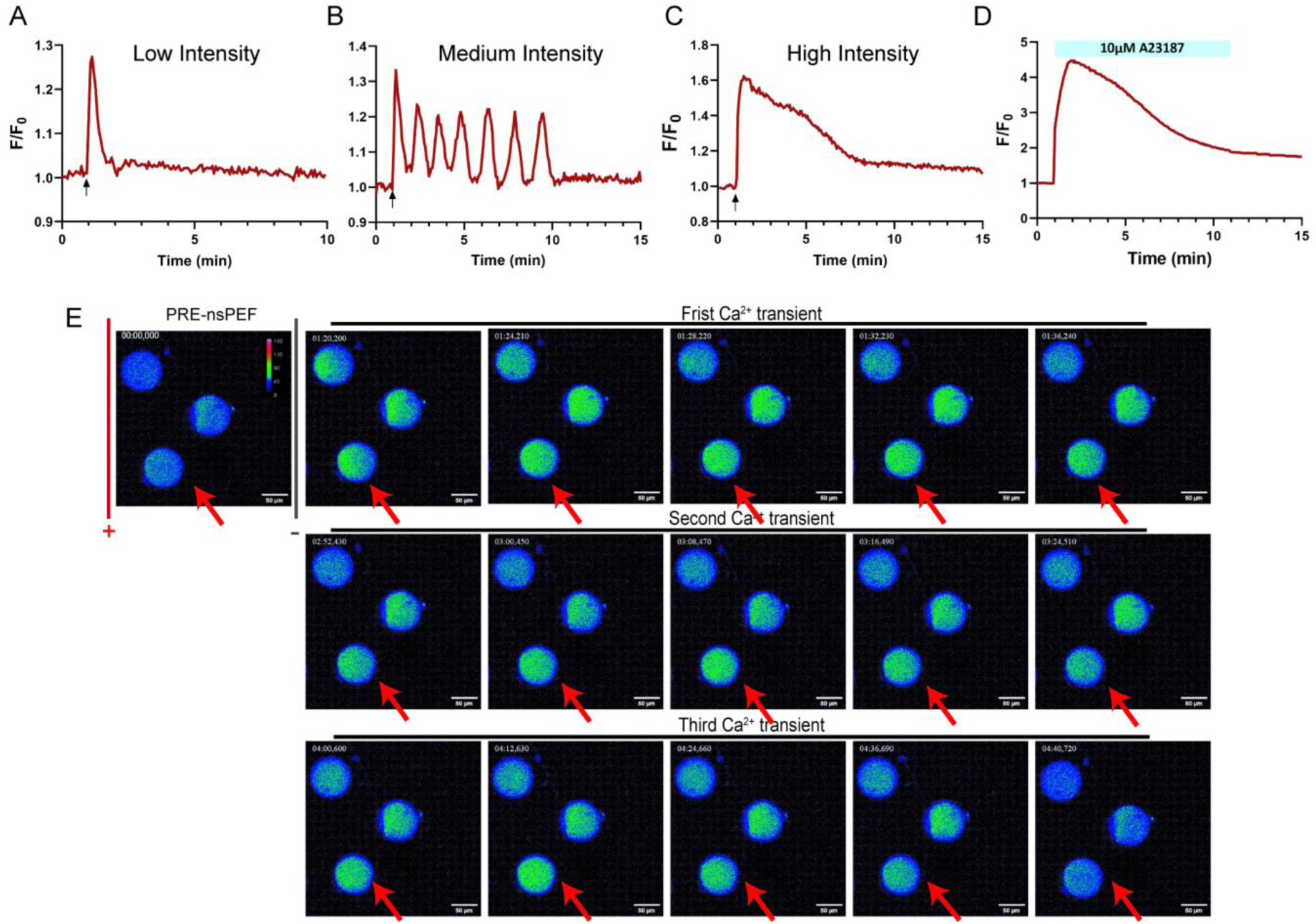
The different patterns of [Ca^2+^]_i_ responses induced by nsPEF stimulation. (A) The pattern of Ca^2+^ response in mouse oocytes stimulated by low-intensity nsPEF (n=26). (B) The pattern of Ca^2+^ oscillations in oocytes stimulated by medium intensity nsPEF (n=24). (C) The pattern of Ca^2+^ response in oocytes stimulated by high intensity nsPEF (n=17). (D) The patterns of Ca^2+^ response in oocytes treated by 10 μM A23187 (n=17). The red line is the representative response from different oocytes in at least three independent experiments (nsPEF pulses—arrows). (E) Representative images of cytoplasmic Ca^2+^ oscillations after nsPEF exposure at medium intensity. The oocyte indicated by red arrow exhibited persistent Ca^2+^ oscillations. Scale bar: 50 μm.

Surprisingly, oocytes exhibited unique spatiotemporal characteristics of [Ca^2+^]_i_ response after moderate-intensity nsPEF stimuli (Figure 1E). Upon pulses of nsPEF, the fluorescence intensity in the ooplasm brightened immediately near the anodal side, followed by swift diffusion of the brighter fluorescence throughout the entire oocyte in few seconds. At this time, [Ca^2+^]_i_ increased rapidly, upregulated by 30.27%, and then returned to its initial state. Subsequently, an unprompted [Ca^2+^]_i_ spikes formed at around 1.35 minutes; meanwhile, the Ca^2+^ level in the whole cytoplasm increased and decreased uniformly. Then, the spontaneous [Ca^2+^]_i_ oscillations recurred in oocytes without additional nsPEF application **(Video 1)**.

The characteristics of the different [Ca^2+^]_i_ responses from each group were set out in **Table 1**. There was no significant difference in the amplitude of [Ca^2+^]_i_ elevation between low intensity group and medium intensity group (*P*=0.4611). The duration and the rise rate of one [Ca^2+^]_i_ transient induced by nsPEF were significantly higher as pulse intensity increased, showed a field intensity dependence in the nsPEF-induced [Ca^2+^]_i_ responses. Moreover, the each parameter of [Ca^2+^]_i_ spike induced by A23187 was significantly higher than counterpart in high intensity group. Collectively, what stands out in Figure 1 is the variability of [Ca^2+^]_i_ response profiles induced by different activation methods.

**Table 1.**
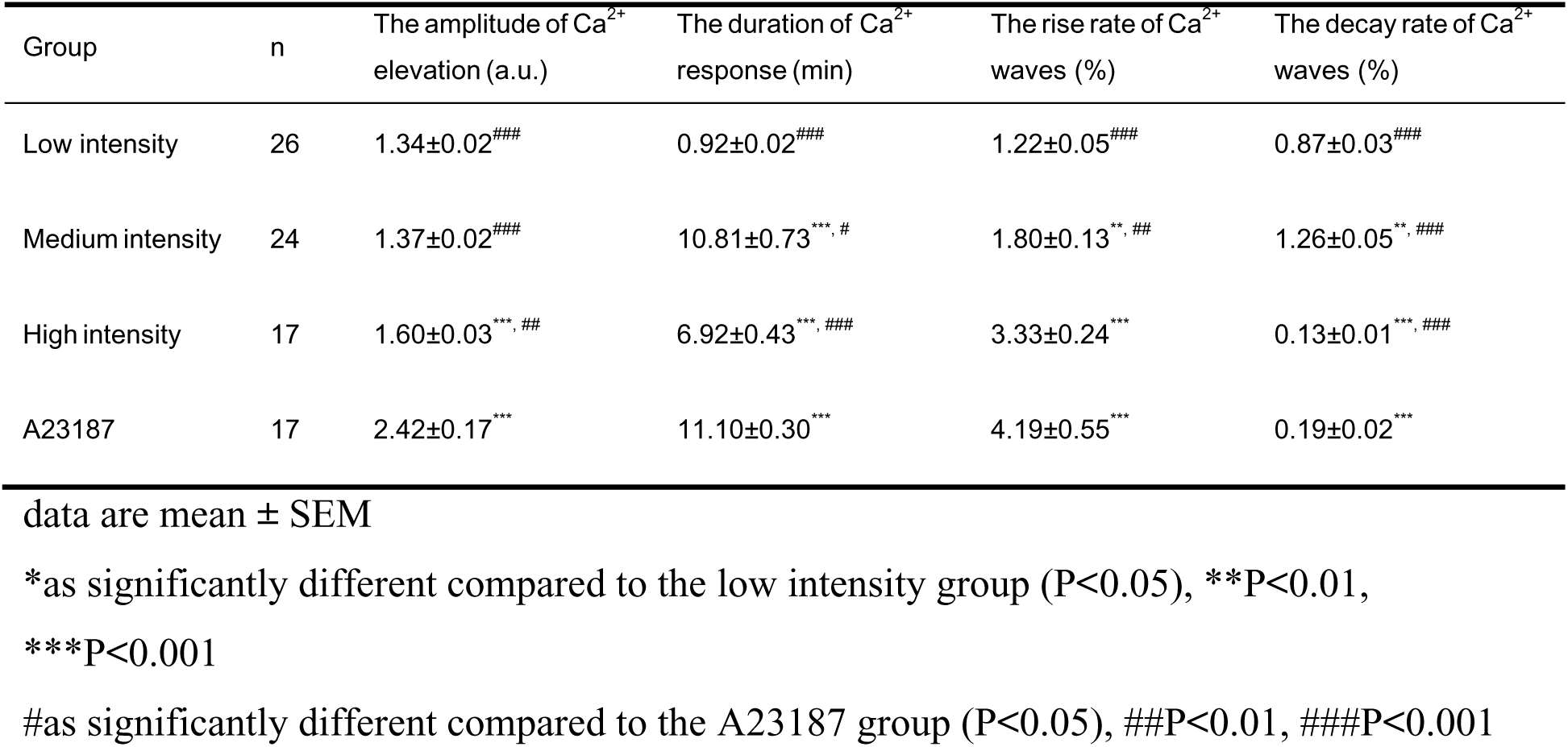
Indications of [Ca^2+^]_i_ responses induced by nsPEF stimulation and A23187.

### 2.2. Electrically controlled Ca^2+^ signaling impacts parthenogenetic activation and preimplantation development in mouse oocytes

We next evaluated whether different pulse-induced Ca^2+^ signaling patterns activate mouse oocyte development in vitro. Oocytes subjected to nsPEF at each field intensity underwent completion of meiosis and pronuclear (PN) formation. The activation rate in medium intensity group was significantly higher than in the other intensity groups of nsPEF stimulation, but not significantly different from that in the A23187 group (81.21% vs 87.27%, *P*=0.8974, **Figure 2A**).

**Figure 2.**
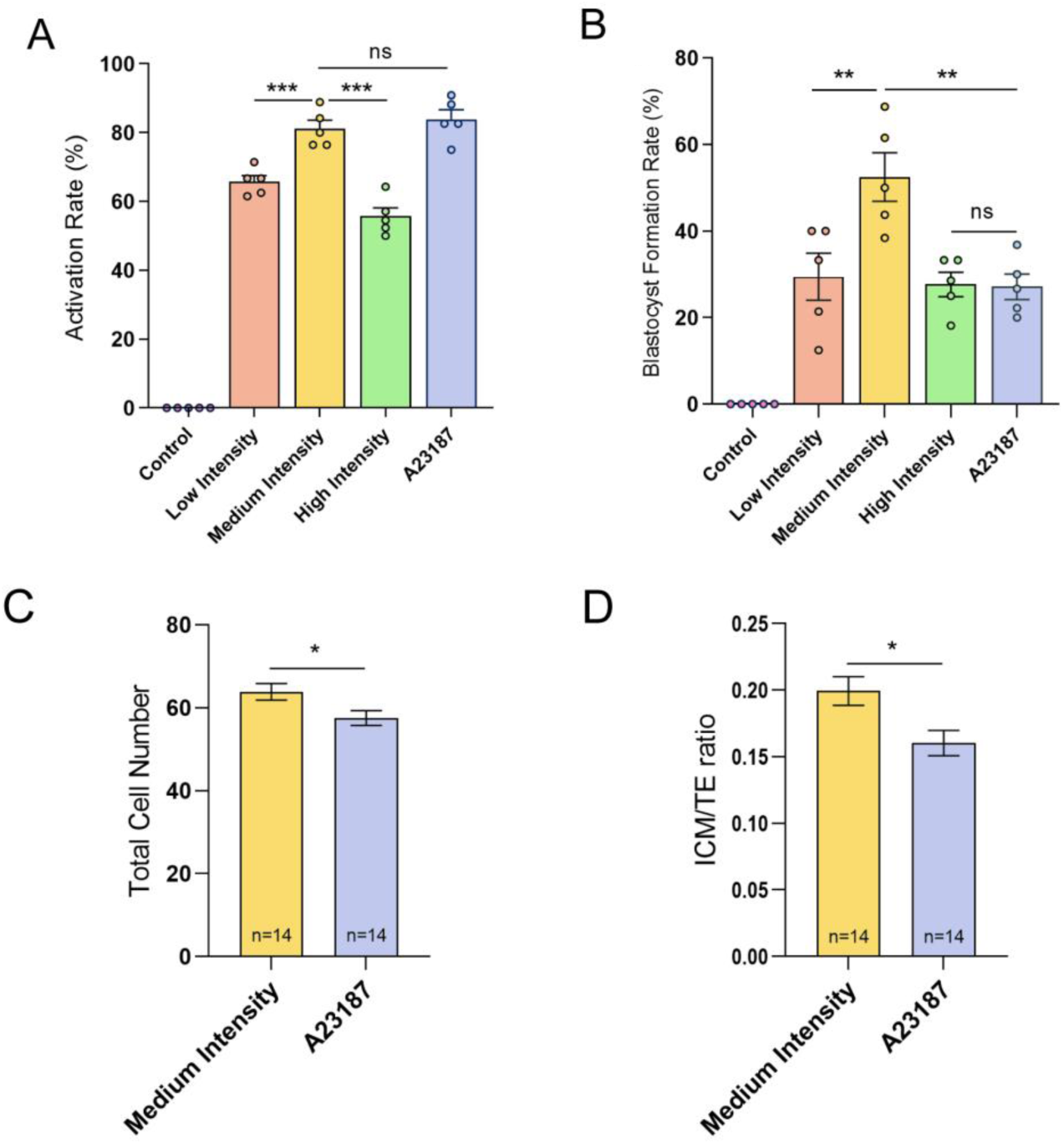
Parthenogenetic activation and developmental outcomes of oocytes after nsPEF stimulation and A23187 treatment. (A) The activation rate of oocytes in different intensity nsPEF treatment groups, A23187 treatment group, and non-stimulation group (control group), respectively. The activation rate is calculated as the ratio of the number of activated oocytes to the total number of treated oocytes. (B) The blastocyst formation rate of oocytes in different groups. This rate is determined by the ratio of developed blastocysts to the total number of activated oocytes. Error bars represent the mean ± SEM of five independent biological replicate experiments (n=5, containing 60, 79, 86, 81, and 84 oocytes in each group respectively) (C) The total cell number of blastocysts in each treatment group. (D) The ICM/TE ratio of blastocysts in each treatment group. The data are presented as mean ± SEM. \**P*<0.05, \*\**P*<0.01, \*\*\**P*<0.001, ns indicates non-significant (*P*>0.05).

Parthenogenetic-activated oocytes from each group further developed to the blastocyst stage, and there were obvious differences in blastocyst formation rates. The blastocyst formation rate in the medium-intensity nsPEFs group was the highest, which was significantly higher than that in the A23187 group (51.67% vs. 26.70%, *P*=0.0016) (Figure 2B), suggesting that medium-intensity nsPEFs improves the developmental potential of activated oocytes compared to A23187 treatment.

Additionally, the development quality of parthenogenetic blastocysts was assessed between medium-intensity nsPEF group and the A23187 group. The parthenogenetic blastocysts in medium-intensity nsPEF group had a higher total cell number (62.43 *vs* 57.5, *P*=0.0258) and a higher ICM/TE ratio (0.20 *vs* 0.16, *P*=0.0275) compared to that in A23187 group (Figure 2C-D), indicating superior embryo quality in medium-intensity nsPEF group.

### 2.3. Fertilization-like Ca^2+^ oscillations simulated by multiple pulses increase oocyte activation efficiency and developmental potentials

The prominent physiological Ca^2+^ changes observed after fertilization is long-lasting and repetitive increases until PN formation. The prolonged time duration of nsPEF-induced Ca^2+^ signaling is probably beneficial for oocyte activation and development in vitro, to this end, the sperm-induced oscillatory Ca^2+^ signals were reproduced in mouse oocytes by repetitive nsPEF pulses. According to the pattern and characteristics of physiological Ca^2+^ oscillation detected during IVF (**Figure 3A** and **Table 2**), oocytes exposed to low-intensity pulses every 3.5 min or medium-intensity pulses every 20 min for 2 hours to reproduce the physiological Ca^2+^ oscillation required for oocyte activation (Figure 3B-C). The comparison of the characteristics of Ca^2+^ oscillation in each group revealed that the [Ca^2+^]_i_ transients elicited by the nsPEF pulses, at either low or moderate intensity, typically exhibit steeper rates of elevation and recovery than natural [Ca^2+^]_i_ spikes (Table 2). In contrast, the spontaneous sustained [Ca^2+^]_i_ oscillations in the oocyte after moderate-intensity nsPEF stimulation more closely resemble the pattern of physiologic [Ca^2+^]_i_ oscillation in IVF group (*P*=0.5819), even though the Ca^2+^ spikes occur at shorter intervals in medium intensity group. Further, compared to single nsPEF stimulation, oocytes exhibited higher parthenogenetic activate efficiency (85.75% and 84.10%, Figure 3D) and blastocyst formation rate (78.13% and 76.28%, Figure 3E) under electro-induced physiological-like [Ca^2+^]_i_ oscillations. Whereas distinctions in [Ca^2+^]_i_ signaling induced by different nsPEF stimuli did not led to differences in oocyte activation and development (*P*=0.7687 and *P*=0.9093), this implies that the Ca^2+^ signaling required for activation may be in a cumulative form instead of strictly dependent on the specific characteristics of the Ca^2+^ spikes.

**Figure 3.**
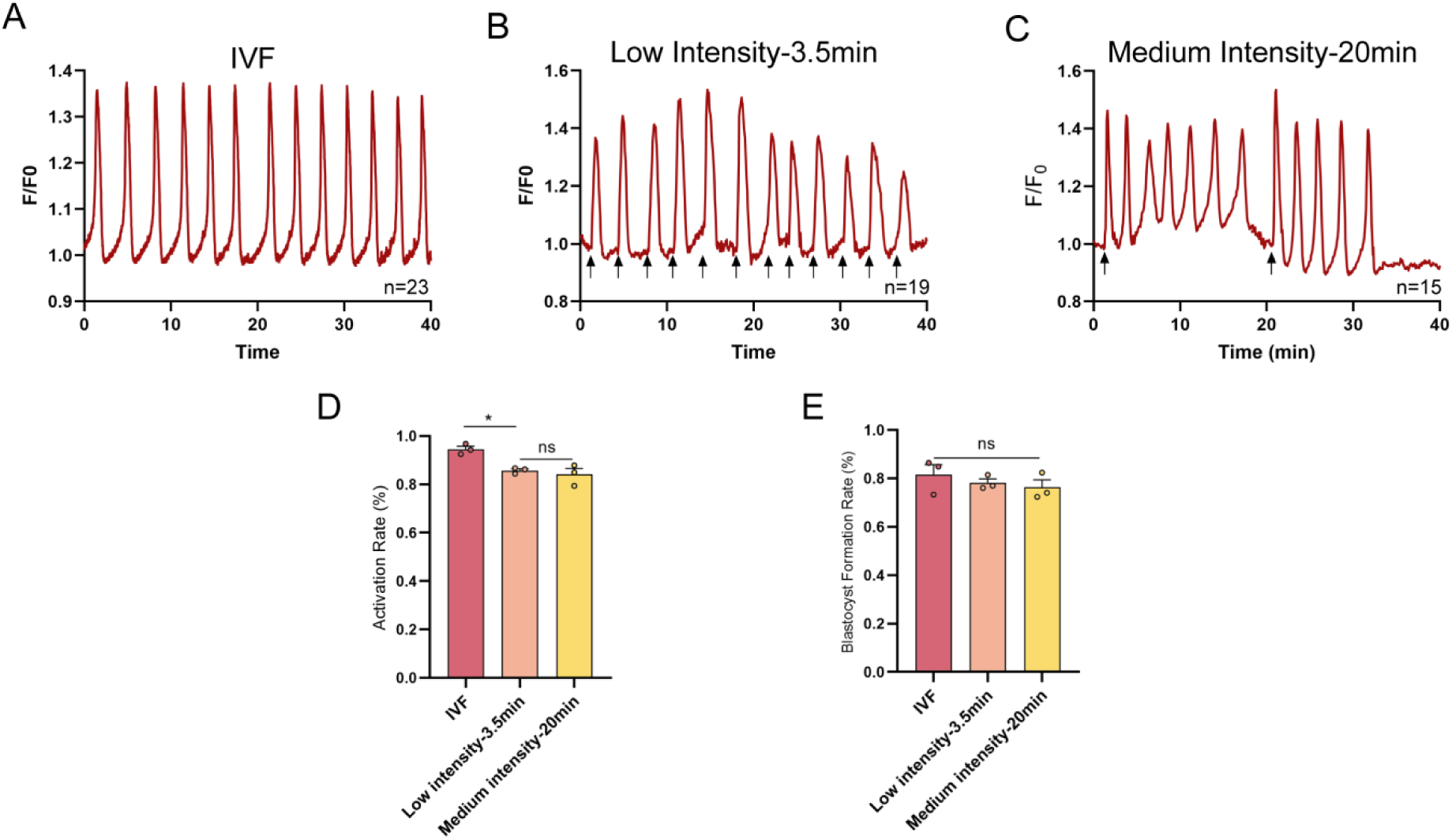
The effect of fertilization-like Ca^2+^ oscillations induced by repetitive nsPEF stimulation on oocyte activation and development (A-C) The different patterns of [Ca^2+^]_i_ oscillations induced by nsPEF stimulation and IVF in mouse oocytes (A) The pattern of Ca^2+^ oscillations in oocytes during IVF. (B) The pattern of Ca^2+^ response stimulated by low intensity nsPEF at 3.5 min intervals. (C) The pattern of Ca^2+^ response stimulated by medium intensity nsPEF at 20 min intervals (nsPEF pulses—arrows). (D) The activation rates of oocytes in different intensity nsPEF treatment groups and IVF group (control group), respectively. (E) The blastocyst formation rates of oocytes in different groups. Error bars represent the mean ± SEM of three independent biological replicate experiments (n=3, containing 106, 91, and 107 oocytes in each group respectively).

**Table 2.**
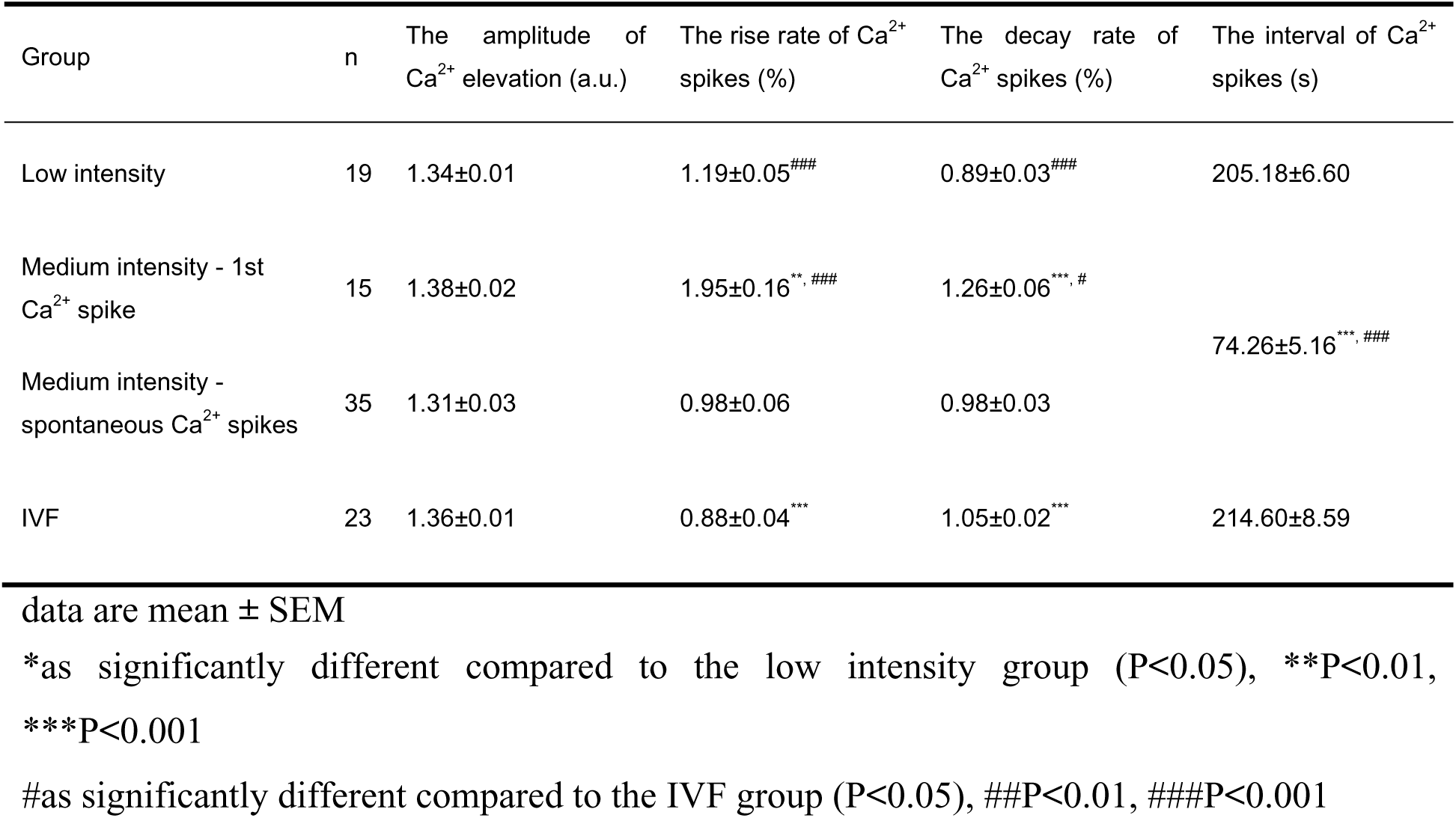
Indications of [Ca^2+^]_i_ responses in different treatment group.

### 2.4. Extracellular Ca^2+^ and intracellular Ca^2+^ storage participate in the formation of [Ca^2+^]_i_ oscillation under nsPEF stimulation

To clarify the formation mechanism of fertilization-like [Ca^2+^]_i_ oscillations elicited by nsPEF pulses, we took advantage of inhibitors to selectively remove free Ca^2+^ from specific regions (**Figure 4A**). First, Ca^2+^ spikes were completely blocked under 0 mM extracellular Ca^2+^ conditions in oocytes treated by EDTA, after low-intensity nsPEFs stimulation (Figure 4B), with the exception that removal of calcium storage from the ER and mitochondria using TG and Ru360 did not compromise the nsPEF-induced Ca^2+^ elevation (Figure 4C). However, it is apparent from Figure 4D that sustained [Ca^2+^]_i_ oscillations could still develop without extracellular Ca^2+^ supplyment after medium-intensity nsPEF stimulation. These oscillatory Ca^2+^ signaling exhibited a lower amplitude (*P*=0.0003), the prolonged duration (*P*=0.0019), and the protracted interval between [Ca^2+^]_i_ transients than those in untreated oocytes (*P*<0.0001, **Table 3**). Therefore, extracellular Ca^2+^ influx is an important component of [Ca^2+^]_i_ oscillation formation induced by nsPEF.

**Figure 4.**
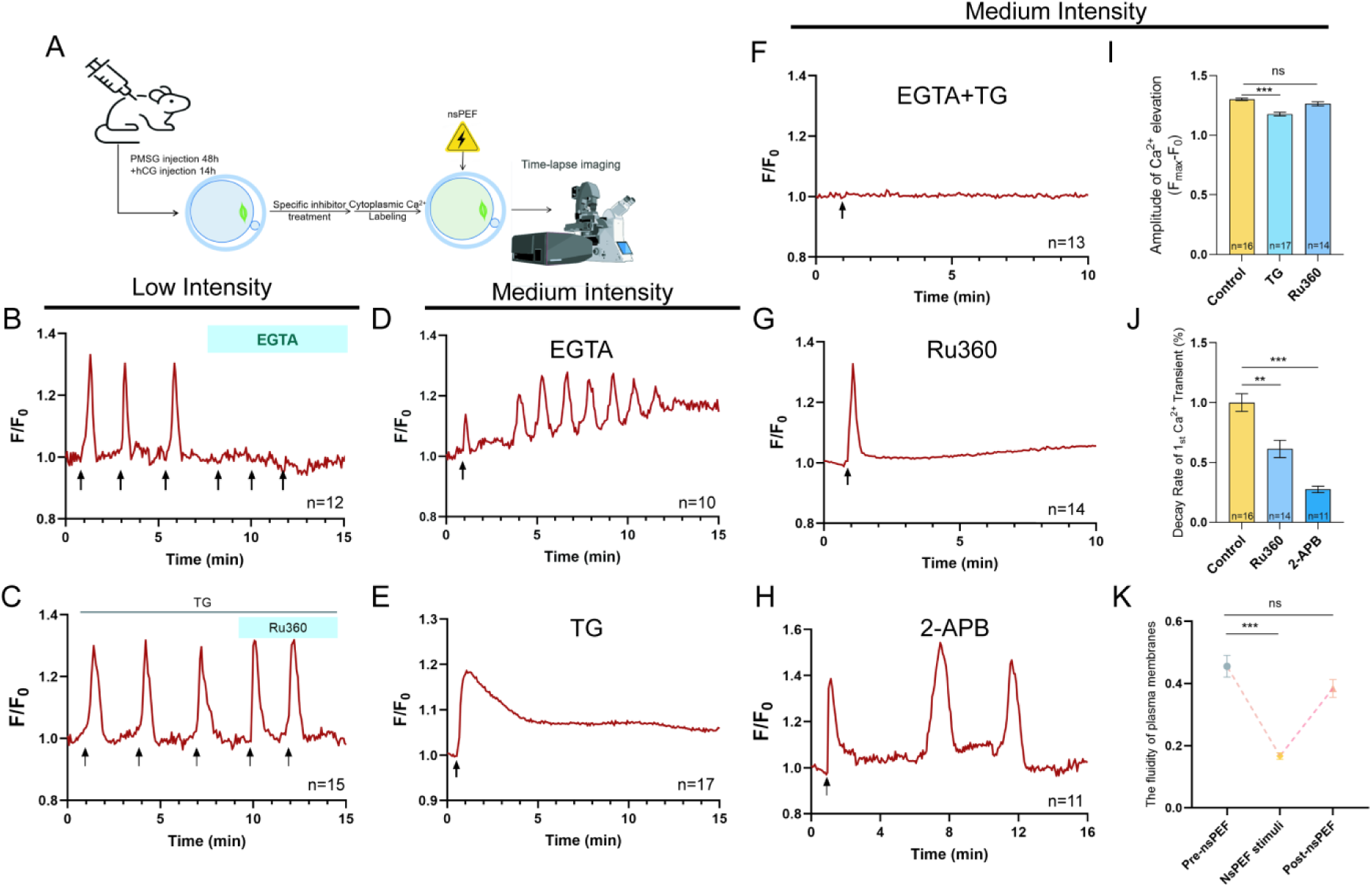
The formation of [Ca^2+^]_i_ oscillations in oocytes upon nsPEF stimulation at low or medium intensity (A) Schematic showing the procedures for the formation pathway of nsPEF-induced [Ca^2+^]_i_ responses by antagonist experiment in mouse MII oocytes. (B) The representative [Ca^2+^]_i_ responses to low-intensity nsPEF stimulation (red line) in oocytes incubated with 5 mM EGTA. (C) The representative [Ca^2+^]_i_ responses to low-intensity nsPEF stimulation (red line) in oocytes incubated with 10 μM TG and/or 10 μM Ru360. (D) The representative [Ca^2+^]_i_ responses to medium-intensity nsPEF stimulation (red line) in oocytes incubated with 5 mM EGTA. (E) The representative [Ca^2+^]_i_ responses to medium-intensity nsPEF stimulation (red line) in oocytes incubated with 10 μM TG (F) The representative [Ca^2+^]_i_ responses to medium-intensity nsPEF stimulation (red line) in oocytes incubated with both 5 mM EGTA and 10 μM TG. (G) The representative [Ca^2+^]_i_ responses to medium-intensity nsPEF stimulation (red line) in oocytes incubated with 10 μM Ru360. (H) The representative [Ca^2+^]_i_ responses to medium-intensity nsPEF stimulation (red line) in oocytes incubated with 10 μM 2-APB (nsPEF pulses—arrows). (I) The amplitude of the first [Ca^2+^]_i_ transient induced by medium-intensity nsPEF from different treatment group. (J) The decay rate of the first [Ca^2+^]_i_ transient induced by medium-intensity nsPEF from different treatment group. (K) The fluidity of plasma membrane on oocytes was detected before stimulation, upon stimulation and after medium-intensity nsPEF stimulation. The data are presented as mean ± SEM. \*\**P*<0.01, \*\*\**P*<0.001, ns indicates nonsignificant (*P* > 0.05).

**Table 3.** Indications of persistent [Ca^2+^]_i_ oscillations induced by nsPEF in medium intensity group.

| Group | n | The amplitude of $Ca^{2+}$ elevation (a.u.) | The duration of intact $Ca^{2+}$ oscillations (min) | The interval of 1st & 2ed $Ca^{2+}$ transient (s) | The number of $Ca^{2+}$ transient |
| --- | --- | --- | --- | --- | --- |
| Control | 16 | 1.33±0.01 | 7.68±0.47 | 1.45±0.10 | 5.94±0.41 |
| EGTA | 10 | 1.13±0.01*** | 10.99±0.47** | 3.30±0.11*** | 5.80±0.57 |
| 2-APB | 11 | 1.23±0.05 | 11.09±0.78** | 6.59±0.85*** | 3.00±0.17*** |

Data are mean ± SEM
\*as significantly different compared to the control group ( $P < 0.05$ ), \*\* $P < 0.01$ , \*\*\* $P < 0.001$

In the absence of Ca^2+^ from the ER storage, oocytes subjected to medium intensity pulses exhibited a single [Ca^2+^]_i_ wave (Figure 4E) with markedly reduced amplitude (*P*<0.0001, Figure 4I), instead of repeated [Ca^2+^]_i_ oscillations. When EGTA and TG were simultaneously employed to clear both extracellular Ca^2+^ and ER stores, there was invisible [Ca^2+^]_i_ alteration after medium intensity nsPEF stimulation (Figure 4F). This suggests that the extracellular milieu and ER stores are the primary contributors to the nsPEF-induced first [Ca^2+^]_i_ transient.

We found that synchronized Ca^2+^ oscillations in cytoplasm and mitochondria were triggered by medium-intensity nsPEF exposure (Figure S1A-B). To verify that mitochondria stores likewise contribute to the nsPEF-induced [Ca^2+^]_i_ oscillations, we specifically depleted Ca^2+^ in mitochondria using 10 μM Ru360 (Figure S1C). In this situation, a single cytoplasmic Ca^2+^ transient was observed in oocytes after nsPEF stimulation (Figure 4G), accompanied by an identical pattern of [Ca^2+^]_mito_ changes (Figure S1D). While the amplitude of [Ca^2+^]_i_ elevation was not significantly altered (Figure 4I, *P*=0.2064), its decay rate was considerably reduced (Figure 4J, *P*=0.0025) in oocytes without mitochondria stores upon nsPEF stimulation. This suggests that mitochondria, as the intracellular fast-response calcium pools, uptake increased [Ca^2+^]_i_ rather than release Ca^2+^ to regulate intracellular calcium homeostasis during the nsPEF-induced [Ca^2+^]_i_ oscillations.

SOCE, as a refilling mechanism for depleted Ca^2+^ stores in ERs, was inhibited by 10 μM 2-APB to determine the role of cytosolic Ca^2+^ backflow into the ER after nsPEF stimulation at medium intensity. The nsPEF-induced [Ca^2+^]_i_ oscillations were significantly attenuated in oocytes (Figure 4H), as evidenced by the prolonged duration of oscillations (*P*=0.0010), the fewer [Ca^2+^]_i_ transients (*P*<0.0001), and the longer intervals between [Ca^2+^]_i_ transients (*P*=0.0008, Table 3). The amplitude of the first Ca^2+^ spiking was comparable (*P*=0.1694, Table 3) but its recovered rate of cytoplasmic Ca^2+^ was significantly lower than that in the untreated group (*P*<0.0001, Figure 4J), demonstrating that the recovery from Ca^2+^ elevation was compromised due to the lack of SOCE function. The results indicated the re-entry of Ca^2+^ into oocytes induced by SOCE was the key point of formation of periodic [Ca^2+^]_i_ oscillation upon medium-intensity pulses.

We further investigated whether modifications to the plasma membrane structure were reversible in oocytes stimulated by nsPEF. The plasma membrane conformation of oocytes was highly responsive to nsPEF stimulation (0.44 vs. 0.17, P<0.0001), and this change in membrane fluidity could be restored within a few minutes (0.44 vs. 0.37, *P*=0.1201), as shown in Figure 4K. This demonstrates that the lifetime of nanopores in the plasma membrane evoked by nsPEF stimuli is limited to only a few minutes, allowing for extracellular Ca^2+^ influx, and then the conformation of plasma membrane rapidly resumes its original state to prevent excessive extracellular Ca^2+^ uptake.

### 2.5. Maintenance of [Ca^2+^]_i_ oscillations is dependent on IP_3_R activation by two regulatory mechanisms

Whether the first [Ca^2+^]_i_ elevation induced by moderate-intensity nsPEF stimulation reaches the activation threshold of the IP_3_R and CICR occurrence to mediate further Ca^2+^ release from the ERs was explored. When IP_3_R activity was inhibited by Xestospongin C (XC), the [Ca^2+^]_i_ oscillation was prematurely halted in oocytes stimulated by medium-intensity nsPEF (**Figure 5A**), indicating that IP_3_R is essential for maintaining nsPEF-induced [Ca^2+^]_i_ oscillations.

**Figure 5.**
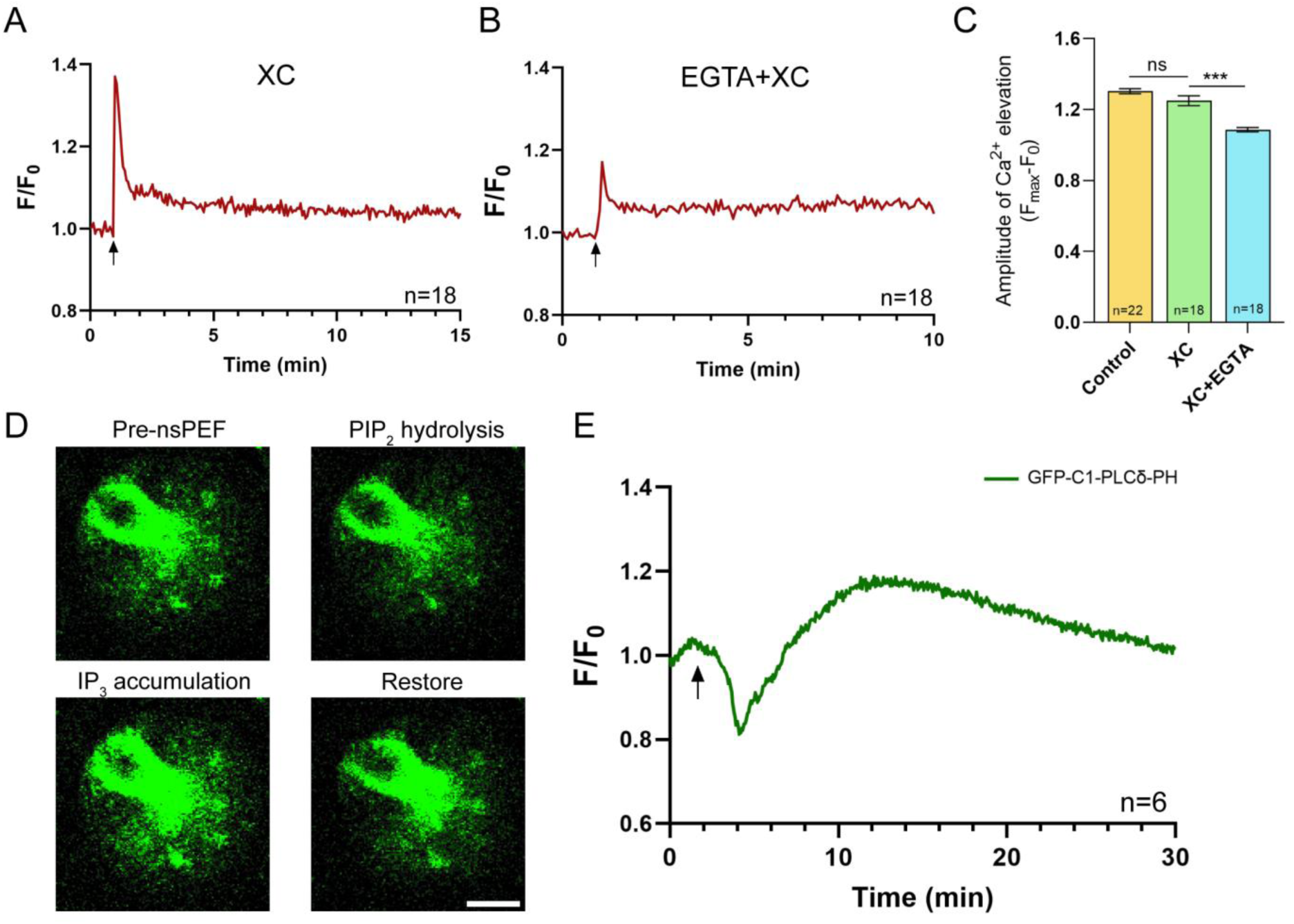
The maintenance mechanism of of Ca^2+^ oscillations stimulated by medium-intensity nsPEF (A) The representative [Ca^2+^]_i_ responses to medium-intensity nsPEF stimulation (red line) in oocytes incubated with 10 μM XC (B) The representative [Ca^2+^]_i_ responses to medium-intensity nsPEF stimulation (red line) in oocytes incubated with both 5 mM EGTA and 10 μM XC. (C) The amplitude of the 1^st^ [Ca^2+^]_i_ transient of [Ca^2+^]_i_ oscillations from different treatment group. (D) Representative images of cytoplasmic GFP-C1-PLCdelta-PH signal alteration. Scale bar: 20 μm. (E) The representative response of GFP-C1-PLCδ-PH fluorescence signal after medium-intensity nsPEF stimulation (green line) in oocytes (nsPEF pulses—arrows). The data are presented as mean ± SEM. \*\*\**P*<0.001, ns indicates nonsignificant (p > 0.05).

There was no significantly difference in the first peak of [Ca^2+^]_i_ increase upon nsPEF stimulation between XC-treated oocytes and untreated oocytes, but oocytes lacking IP_3_R activity exhibited a significantly reduced Ca^2+^ peak in 0 mM extracellular Ca^2+^ milieu (Figure 5B-C). This suggests that nsPEF pulses did not directly trigger Ca^2+^ efflux across IP_3_R channels on ERs. The Ca^2+^ elevation evoked by nsPEF secondarily activates IP_3_R to form spontaneous [Ca^2+^]_i_ oscillations via the CICR pathway.

Based on the model of quantitative regulation of intracytoplasmic Ca^2+^-IP_3_ kinetics by nsPEF ^31^, we speculate that the generation of IP_3_ similarly activated IP_3_R to maintain [Ca^2+^]_i_ oscillations after nsPEF stimulation. Pre-exposure GFP-C1-PLCδ-PH fluorescence in oocytes illustrated the presence of intracellular PIP_2_ in vesicular structures (Figure 5D). The green fluorescence decreased rapidly at 2.71±0.30 minutes after nsPEF exposure, then beginning to gradually recover at 3.44±0.27 minutes (Figure 5D-E), implying that nsPEF stimulation promotes intracellular PIP_2_ hydrolysis and increases IP_3_ concentration in oocytes. Under the dual regulation of Ca^2+^ level and IP_3_, the activated IP_3_R mediates Ca^2+^ release from ERs to generate sustained [Ca^2+^]_i_ oscillations.

### 2.6. The application of nsPEF stimulation is a safe method for oocyte activation

The safety of nsPEF applications on mouse oocytes was further examined. The early apoptosis rates of oocytes in the low to high intensity nsPEF groups were not significantly different from those in the control (non-stimulation) group; despite the apoptosis rate in the A23187 groups was highest, all rates remained below 20% (**Figure 6A**). Detection of intracellular ROS and GSH concentrations in oocytes from each group revealed greater ROS accumulation and GSH depletion with increasing field intensity of nsPEF exposure. However, the level of oxidative stress was significantly higher in A23187-treated oocytes than that in high intensity group (Figure 6B-C).

**Figure 6.**
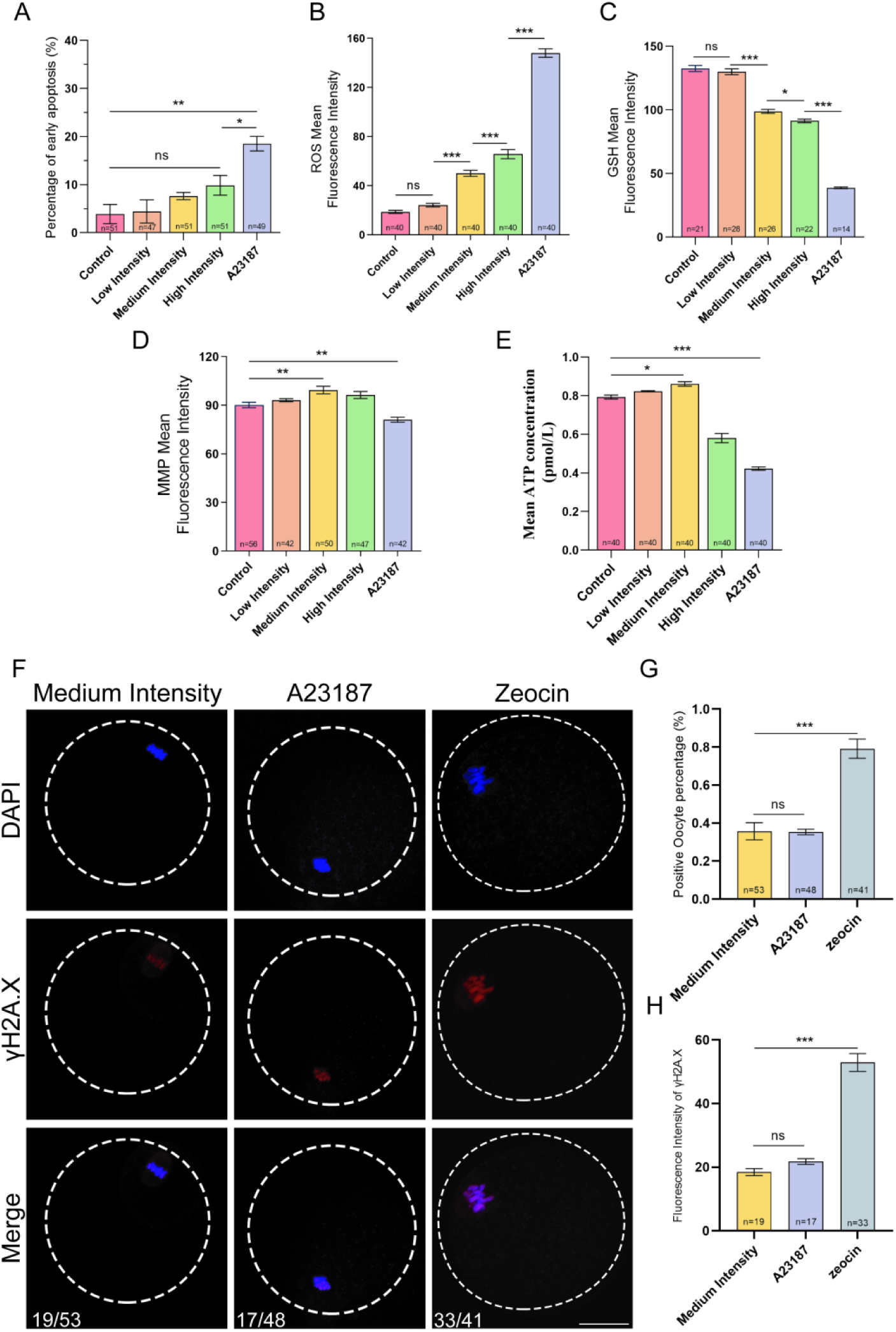
The effects of nsPEF stimulation on cytoplasmic and nucleus security in mouse oocytes (A) Percentage of early apoptosis of oocyte in each treatment group. (B) Mean fluorescence intensity of intracellular ROS in each treatment group. (C) Mean fluorescence intensity of intracellular GSH in each treatment group. (D) Mean fluorescence intensity of MMP in oocytes in each treatment group. (E) Intracellular ATP content of each oocyte in each treatment group. (F) Representative immunostaining images of oocytes with DNA double-strand breaks in each treatment group. Scale bar: 20 μm. (G) The percentage of oocytes with DNA damage in each treatment group. (H) The fluorescence intensity of oocytes with DNA damage in each treatment group. The data are presented as means ± SEMs. \**P*<0.05, \*\**P*<0.01, \*\*\**P*<0.001, ns indicates nonsignificant (*P* > 0.05).

Notably, significantly higher mitochondrial membrane potential (MMP) levels (*P*=0.0023) and greater ATP concentration (*P*=0.0341) were observed in oocytes from the medium-intensity group compared to the control group (Figure 6D-E), suggesting that oocytes stimulated by medium-intensity nsPEF had higher mitochondrial activity. Conversely, A23187 treatment caused lower MMP (*P*=0.0049) and reduced ATP content in oocytes (*P*<0.0001).

The potential impairment of DNA structure across the nuclear envelope by nsPEF pulses was investigated using γH2A.X antibody labeling (Figure 6F). Both nsPEF exposure and A23187 incubation resulted in a lower incidence of abnormal chromosome distribution and DNA double-strand breaks (Figure 6G) and a lower degree of DNA damage (Figure 6H) compared to zeocin-treated oocytes (positive control). This suggests that nsPEF stimulation in this protocol is non-damaging to DNA in oocytes.

### 2.7. Human oocytes exhibit various degrees of Ca^2+^ waves after different parameter of nsPEF stimulation

To investigate how [Ca^2+^]_i_ changes in human oocytes under nsPEF exposure, we found that a single Ca^2+^ transient was induced in ooplasm when different intensities of nsPEF were applied to human IVM-MII oocytes. Oocytes exhibited a single, minimally visible [Ca^2+^]_i_ wave when stimulated by 10 pulses of low-intensity nsPEF, while the amplitude of [Ca^2+^]_i_ wave increased after medium and high intensities stimulation (**Figure 7A-B**). Oocytes also formed increasing [Ca^2+^]_i_ waves under cumulative pulses of medium-intensity nsPEF (Figure 7C-D). Cytoplasmic Ca^2+^ concentration was urgently upregulated by 34.84% under 30 pulses of medium-intensity nsPEF. Thus, varying degrees of Ca^2+^ peaks could be induced by adjusting the intensity or number of nsPEF pulses.

**Figure 7.**
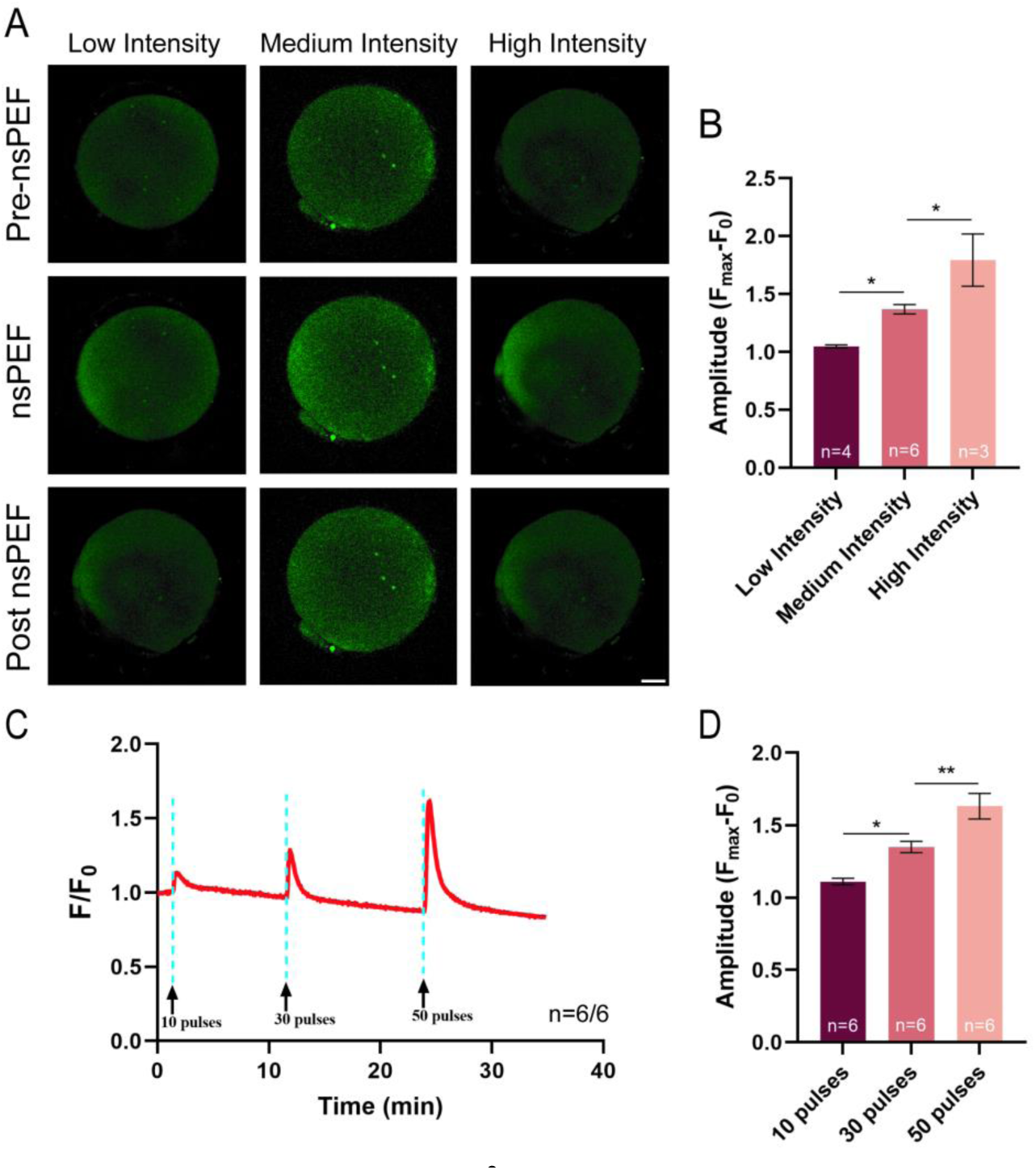
The patterns of cytoplasmic Ca^2+^ waves in human oocytes induced by nsPEF stimulation (A) Representative images of cytoplasmic Ca^2+^ alteration before and after nsPEF exposure of 10 pulses at low, medium or high intensity. Scale bar: 20 μm. (B) The amplitude of Ca^2+^ transient induced by nsPEF with different intensities. (C) Representative Ca^2+^ oscillation patterns following repeated nsPEF exposure with medium intensity. (D) The amplitude of Ca^2+^ transient induced by nsPEF with different number of pulses. The data are presented as mean ± SEM. \**P*<0.05, \*\**P*<0.01.

## 3. Discussion

Patients with OAD face physical and psychological damage from repeated ICSI cycles, but this obstacle has not been fully overcome by current AOA methods and their processes remain unclear. Therefore, there is an urgent need for the revolutionary and improved AOA strategies. In this study, we propose an electrical means of AOA using nsPEF technology, designed to ensure the survival and safety of oocytes while enhancing oocyte activation efficacy and subsequent early embryonic development by generating fertilization-like propagating Ca^2+^ oscillations *in vitro*. NsPEFs stimulation at the low intensity induces extracellular Ca^2+^ influx, whereas, moderate-intensity nsPEF pulses prompt external Ca^2+^ influx and Ca^2+^ release from organelles in oocytes, followed by cytoplasmic Ca^2+^ elevation to induce intracellular PIP_2_ hydrolysis, regulating IP_3_R-related intrinsic calcium homeostasis, which are involved in maintaining sustained Ca^2+^ oscillations. Thus, intracellular physiological-like Ca^2+^ oscillations are formed during oocyte activation by repeated nsPEF stimulation. This approach is different from prior research by replacing chemical agents with a physics-driven, non-invasive technique that preserves oocyte structural integrity and enhances developmental potential. Consequently, the development of the nsPEF technique holds promise as an innovative and adaptable approach for refining AOA treatments.

Most AOA methods can only induce a single or a limited number of Ca^2+^ transients in the ooplasm, failing to replicate the physiological pattern of fertilization-induced Ca^2+^ oscillations. This limitation results in suboptimal activation and poor embryonic developmental competence. Consequently, optimizing AOA technology has focused on mimicking the physiological Ca^2+^ oscillations at fertilization in oocytes through non-invasive means. In this study, we manipulated [Ca^2+^]_i_ levels in mature oocytes using nsPEF with minimal electro-damage, exploring the correlation between the pattern of cytoplasmic Ca^2+^ changes and the oocyte activation outcomes. Different patterns of cytoplasmic Ca^2+^ changes were induced by nsPEF stimulation at varying intensities. Low-intensity pulses caused rapid [Ca^2+^]_i_ elevation followed by acute restoration. Repeated low-intensity nsPEF stimulations successfully generated fertilization-like Ca^2+^ oscillations. As a result of these oscillations, oocyte activation efficiency increased from 66% to 86%, highlighting the importance of the duration of [Ca^2+^]_i_ oscillations. Interestingly, the sustained [Ca^2+^]_i_ oscillation was triggered by individual moderate-intensity electrical pulses, consisting of 5 to 8 Ca^2+^ spikes, accompanied by mitochondrial Ca^2+^ oscillation. This unique oscillatory pattern led to a higher number of oocytes developing into blastocysts compared to other nsPEF pulse intensities. Furthermore, extending the duration of these physiological-like [Ca^2+^]_i_ oscillations by repeated nsPEF applications significantly increased the number of both activated oocytes and developed blastocysts.

In contrast, with increasing nsPEF pulse intensity, a continuous cytoplasmic Ca^2+^ elevation was observed in the oocytes. This pattern of Ca^2+^ signaling differed from the brief Ca^2+^ spikes previously described and was more comparable to the broad Ca^2+^ waves induced by A23187. More oocytes treated with A23187 resumed meiosis and formed PN compared to those exposed to high-intensity nsPEFs. This difference is likely due to stronger Ca^2+^ signal after A23187 treatment, which sufficiently inhibits maturation-promoting factor (MPF) activity, thus activating the oocytes. However, this non-physiological Ca^2+^ wave resulted in lower blastocyst formation rate. Oocytes activated by electrical pulses typically developed to the blastocyst stage and differentiated into ICM and TE. Both indicators of blastocyst quality were significantly enhanced in oocytes subjected to medium-intensity electrical stimulation, suggesting superior blastocyst quality compared to those in the A23187-treated group.

Furthermore, several studies have shown that electrical activation significantly improves both the developmental rate and quality of blastocysts ^32,33^, supporting the feasibility and effectiveness of nsPEF for oocyte activation.

Although our findings indicate that oocytes exposed to constant high [Ca^2+^]_i_ remain universally activated, oscillating Ca^2+^ signals during activation significantly improved embryonic development. Prolonging the duration of these [Ca^2+^]_i_ oscillations increases the number of oocytes that respond to activation signals and successfully progress to the preimplantation stage in vitro. Despite certain discrepancies between the form of electrically induced [Ca^2+^]_i_ oscillations and physiological conditions, these did not impact the rates of oocyte activation and development, implying that oocytes can accommodate a range of Ca^2+^ oscillatory parameters. Single or oscillatory Ca^2+^ signals could be interpreted by oocytes to execute diverse biological processes necessary for activation ^34^. This combination of prolonged duration and oscillatory patterns may reflect an evolutionary strategy in mammals. Sufficient Ca^2+^ signaling activates Ca^2+^-sensitive kinase CaMKII, leading to APC activation, cyclin B degradation, and downregulation of MPF activity, which are essential for MII exit and thorough meiotic cycle completion ^10^. Although the role of oscillatory Ca^2+^ signals in early embryonic development was still unclear, Ozil et al. demonstrated that the absence of these Ca^2+^ signals at fertilization affects gene expression in blastocysts and decreases the rate of preimplantation ^35^. Ca^2+^ oscillations are assumed to be a crucial signal during oocyte activation, exerting broad and profound effects on various critical aspects of fertilized oocyte development.

Considering that physiological-like Ca^2+^ responses are formed upon stimulation with low-or moderate-intensity nsPEF, it is necessary to probe how [Ca^2+^]_i_ evolution induced by nsPEF stimulation. On the basis of a site-by-site exclusion of cytoplasmic Ca^2+^ sources, the Ca^2+^ elevation evoked by low intensity nsPEF was shown to only be dependent on extracellular Ca^2+^ influx. The [Ca^2+^]_i_ elevation upon nsPEF stimulation at moderate intensity involves contributions from extracellular medium and the ER, evidenced by disruptions in [Ca^2+^]_i_ oscillations patterns in oocytes pretreated with inhibitors such as EGTA and TG. Subsequently, the increased Ca^2+^ in the cytoplasm is quickly restored to its initial state by the backflow into ERs and mitochondria. Oocytes pretreated with Ru360 failed to pump Ca^2+^ into mitochondria, disrupting the original intracellular calcium regulatory homeostasis and leading to the premature interruption of nsPEF-induced [Ca^2+^]_i_ oscillations. The formation of spontaneous [Ca^2+^]_i_ oscillations was also delayed by inhibiting SOCE which impedes the refilling of the ER storage. This indicates that both ERs and mitochondria are involved in recycling cytoplasmic Ca^2+^ and secondary replenishment during [Ca^2+^]_i_ oscillation. The nanopores in the plasma membrane stimulated by nsPEF were rapidly restored as before, as demonstrated by the cell membrane fluidity assay. Taken together, in response to nsPEF stimulation, the plasma membrane and membranes of ER were charged to form transient nanopores mediating free Ca^2+^ circulation in each compartment, followed by restoration of intracellular calcium homeostasis in dependence on several Ca^2+^ regulatory tools.

The mechanism of spontaneous formation of [Ca^2+^]_i_ oscillations after nsPEF stimulation was further explored. Inhibition of IP_3_R activity leads to an early cessation of propagating Ca^2+^ oscillations by disrupting CICR mechanism. Furthermore, the nsPEF stimulation also leads to the downregulation of PIP_2_ and subsequent production of IP_3_ in oocytes. This supports that nsPEF pulses activate the PIP_2_-IP_3_-IP_3_R pathway and regulate intracellular calcium reservoirs to generate a physiological pattern of [Ca^2+^]_i_ oscillation. The biphasic dependence of Ca^2+^ channel IP_3_R gives rise to [Ca^2+^]_i_ oscillations, whose structural properties exhibit rapid positive feedback and slow negative feedback processes on cytosolic Ca^2+^. ^31^ It’s possible that PIP_2_ hydrolysis and IP_3_ generation triggers Ca^2+^ release in the ERs, which explains why oocytes remain able to generate [Ca^2+^]_i_ oscillations in the absence of external Ca^2+^ (Figure 4 D) or SOCE activity (Figure 4 H). Additionally, the simulation model of nsPEF-induced [Ca^2+^]_i_ oscillations detailedly explains that the Ca^2+^ increase significantly activates IP_3_R activity at moderate intensity pulses.^31^ We revealed that nsPEF stimulation induced both cytoplasmic Ca^2+^ increase and IP_3_ production to activate the IP_3_R, which is engaged in the maintenance of [Ca^2+^]_i_ oscillations. Under physiological conditions, fertilization-induced Ca^2+^ oscillations are mostly dependent on IP_3_R-mediated Ca^2+^ release from the ER.^36^ The generation of IP_3_ through PLC-mediated hydrolysis of PIP_2_ is crucial for Ca^2+^ oscillation during fertilization across various species.^37–39^ It is reasonable to speculate that nsPEF stimulation induces the generation of [Ca^2+^]_i_ oscillations in a physiological manner to increase oocyte activation and developmental potential.

At the onset of fertilization in mammals, a cortical flash and wave-like oscillations first appear at the site of sperm-oocyte interaction, rapidly causing the [Ca^2+^]_i_ increase and spreading across the oocyte.^36,40^ Notably, oocytes exhibit similar spatiotemporal properties when exposure to medium-intensity nsPEF (shown in Figure 1 E). Calcium ionophores A23187 transport Ca^2+^ across cell membranes and induce extracellular Ca^2+^ influx by increasing their permeability.^41^ This results in a steady Ca^2+^ increase, forming the single Ca^2+^ transient (shown in Figure 1 D). Although the Ca^2+^ peak induced by A23187 exceeds the threshold required for oocyte activation, it causes suboptimal activation and diminished developmental potential due to the non-physiological patterns of Ca^2+^ release.

In addition to analyzing [Ca^2+^]_i_ oscillatory patterns, the safety of nsPEF exposure on oocytes was evaluated. The incidence of oocyte apoptosis and oxidative stress increased positively with nsPEF intensity. Oocytes exposed to A23187 exhibited higher levels of oxidative stress and apoptosis, indicating that nsPEF are a safer method for in vitro activation compared to A23187. Previous studies support our conclusions.^42,43^ Notably, oocytes stimulated by moderate-intensity nsPEF showed more active mitochondrial function. The charging effect of nsPEF stimulation appears to affect the mitochondrial respiratory chain, promoting ATP production. ^44,45^ However, stronger evidence is required from further exploration. In contrast, the large Ca^2+^ influx mediated by A23187 triggered oxidative stress and decrease mitochondrial activity, suggesting that this non-physiologic Ca^2+^ wave impairs the cytoplasmic quality of oocytes. It has been confirmed that intense electrical pulses can exceed the charging time constant of the nuclear membrane, increasing its conductivity and forming nanopores.^46^ The 10-ns electrical stimulation used in our protocols did not directly cause significant DNA double-strand breaks or aberrant chromatin alignment, and the levels of DNA damage were not statistically different from those in the A23187 treatment group.

Therefore, the genetic material disruption within the oocyte is not a concern for either A23187 treatment or nsPEF stimulation. To ensure the security and effectiveness of nsPEF in AOA, employing a controlled 10-ns PEF setting allows for a broader safety margin without adverse effects.

Numerous AOA problems are associated with the activation of unfertilized oocytes by A23187: a single and excessive elevation of [Ca^2+^]_i_ concentration, non-physiological Ca^2+^ transport, and oxidative damage to cellular activity. ^47^ However, A23187 remains the most commonly employed AOA strategy in clinical practice ^12,13^. Therefore, the development of more secure and efficient AOA technology is urgently needed. Application of a single nsPEF stimulation induced a significant Ca^2+^ spiking in human oocytes, with the amplitude of its peak varying with stimulus intensity and the number of pulses. Repeated nsPEF stimulation induces Ca^2+^ oscillations in human oocytes, and the characteristics of this [Ca^2+^]_i_ transient closely resemble the [Ca^2+^]_i_ waves observed during fertilization (Figure 7 C). We speculate that [Ca^2+^]_i_ oscillations during human fertilization can be accurately replicated by applying specific quantities of nsPEF pulses at precise intervals, thereby facilitating meiosis completion and enhancing developmental potential. Therefore, it is expected that applying nsPEF pulses several times can mimic sperm-activated [Ca^2+^]_i_ oscillations in unfertilized oocytes. This physiological-like [Ca^2+^]_i_ oscillation induced by nsPEF stimulation could overcome the activation and developmental defects of current AOA methods.

However, nanopores in cell membranes stimulated by nsPEF cannot be directly observed due to experimental instrument limitations. The activation of the intracellular PIP_2_-IP_3_-IP_3_R pathway after nsPEF stimulation needs to be verified by signaling cascade experiments, which is our future research objective. The disparities in the intensity of nsPEF exposure on human and mouse oocytes are possibly linked to different cell diameters. The intensity of the pulse that penetrates the cell membrane of a spherical cell increases as its diameter increases. ^48^ Different nsPEF pulse parameters are required to produce intracellular effects in different cell types ^49^, the electrical parameters of nsPEF suitable for human oocyte activation remain to be further determined. Despite the limited quantity and quality of collected human oocytes, nsPEF stimulation still accurately modulated intracellular Ca^2+^ changes, suggesting its potential to induce a physiological-like [Ca^2+^]_i_ oscillation pattern in unfertilized oocytes. We will next collect a large number of human unfertilized oocytes to determine appropriate nsPEF stimulation conditions, thereby supporting the argument that nsPEF stimulation is a relatively better means of AOA at present.

In summary, we systematically explored the relationship between [Ca^2+^]_i_ signaling profile finely modulated by nsPEF pulse intensity, and oocyte activation and development. Notably, nsPEF pulses with moderate intensity mobilize the intracellular calcium stores to generate a unique pattern of cytoplasmic Ca^2+^ oscillations in a physiological manner, which promotes the completion of meiosis and early embryonic development in mouse oocytes. The electrical intensity threshold of nsPEF stimulation regulating Ca^2+^ changes in human oocytes has also been revealed. The multiple Ca^2+^ spike patterns induced by nsPEF suggest that this technique could potentially be applied clinically to enhance fertilization success in patients experiencing recurrent total or near-total ICSI failure due to sperm activation deficits.

## 4. Experimental Section/Methods

*Ethics Statement:* This study was registered at Medical Ethics Committee of Peking University People’s Hospital (PKUPH) prior to inclusion (No. 2020PHB397-01). All procedures involving live animal handling and euthanasia were approved by the Institutional Animal Care and Use Committee & Ethic Review Committee of PKUPH (No. 2020PHE099) and the Ethic Review Committee of Beijing Chao-Yang Hospital, Capital Medical University (No. 2022-KE-436). Unless otherwise stated, all chemicals and medicines were purchased from the Sigma Chemical Co. (St Louis).

*Animal experiments:* Seven-week-old CD-1® (ICR) mice were acquired from the Beijing Vital River Laboratory Animal Co., Ltd., and housed under standard conditions in the animal experimentation facility. These mice were maintained in a pathogen-free environment with 12-hour light/dark cycles for two weeks.

*Mouse oocytes collection and culture in vitro:* Mouse mature oocytes were harvested from 9-week-old female ICR mice following ovulation induction (*n*=9 per group), which involved the intraperitoneal injection of 10 IU of pregnant mare serum gonadotropin (PMSG), followed 46-48 hours later by an injection of 10 IU of human chorionic gonadotropin (hCG), both from Ningbo Second Hormone Factory. MII stage oocytes were released from the oviducts at 14 h post-hCG administration and cumulus cells were dissociated using 0.1% (w/v) hyaluronidase. The cumulus-free oocytes were then washed by three times and maintained in M2 medium under mineral oil at 37°C and 5% CO_2_ until further experimental procedures.

*Nanosecond pulse exposure on mouse oocytes:* Oocytes were collected and randomly grouped, then they were stimulated by low, medium or high intensity nsPEF. A custom-built nsPEF generator, delivering pulses of 10 ns duration, was connected to a petri dish with parallel electrodes to applying pulses on the oocytes (Figure S2A-B). Oocytes were subjected to 10 pulses of nsPEF at different indensity, each with a duration of 10 ns. The intervals between pulses were approximately 1 second.

Considering the different volumes of the M16 medium used in the self-made electrode dish (Figure S2C) versus the platinum Petri dish (Harvard Bioscience, MA), we established a relationship between the electric pulses in different electrode dishes based on the electrical doses of nsPEF. The electrical dose of nsPEFs is chiefly determined by pulse intensity, field duration, and pulse count, while the resistance of medium between parallel electrodes also influences the electrical dose exposed by oocytes. The formula for calculating the electrical dose between parallel electrodes is as follows:

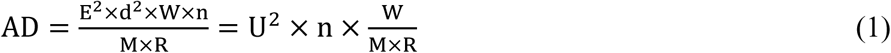

Here, AD represents the absorbed dose (J/g), which indicates the electrical dose from nsPEFs stimuli; E is the applied nsPEFs’ pulse intensity (V/m); d stands for the pulse electrode gap (m); W signifies the pulse width (s); n is the number of nsPEFs pulses applied; R depicts the resistance of the culture medium, and M represents the volume of solution between the parallel electrodes. Consequently, the relationship between the AD in different stimulation dishes is exhibited in Table S1.

Specifically, the low intensity group manifested the electric dose of 7-8 kV/cm applied to the platinum dish was equivalent to the electric dose of 16-17 kV/cm in the self-made confocol dish; the medium intensity group manifested the equal electric dose of 9.5-10.5 kV/cm in the platinum dish and 20-21 kV/cm in the self-made confocol dish; the high intensity group manifested the equal electric dose of 12.5-14 kV/cm in the platinum dish and 24-25 kV/cm in the self-made confocol dish.

*Mouse oocyte activation and in vitro culture:* Following exposure to different nsPEF pulses, oocytes were thoroughly washed and cultured in KSOM medium under mineral oil at 37°C in a humidified atmosphere containing 5% CO2 during the embryonic development. Oocytes were incubated in M16 medium containing 10 μM A23187 for 10 min at 37°C and 5% CO_2_ as a positive control, and in vitro culture in same protocol. Oocytes placed in Petri dishes but exposed to 0 kV/cm pulses served as the sham exposure group (control group). Oocytes that form 1-2 pronuclei or cleave were considered as activated oocytes, and they were cultured in KSOM medium and tracked at 24-hour intervals for up to 5.5 days to monitor embryo development.

*Measurement of intracellular ROS, glutathione (GSH), and oocytes apoptosis*: Intracellular reactive oxygen species (ROS) or glutathione (GSH) levels in oocytes were detected using 1 mmol/L 2’,7’-dichlorofluorescein diacetate (DCFH-DA) or 10 μmol/L Cell Tracker Blue (Invitrogen, Carlsbad, CA). Images were captured using a fluorescence microscope (Olympus IX73) and quantified with EZ-C1 Free-Viewer software (Nikon, Tokyo, Japan). To assess early apoptosis in oocytes, staining oocytes was performed with the Annexin V-FITC/PI Apoptosis Detection Kit (Vazyme, Nanjing, China), following the manufacturer’s protocol. Oocytes were incubated with 5 µl of Annexin V-FITC and 5 µl of Propidium Iodide (PI) for 10 minutes in the dark at 37°C, then thoroughly washed three times with PBS. Green fluorescence localized to the zona pellucida indicated non-apoptotic cells, while green fluorescence present on both the membrane and zona pellucida signaled apoptosis. PI can penetrate the cell membrane of necrotic or late apoptotic cells, staining the nucleus to red fluorescence. The fluorescence signals were visualized using a fluorescence microscope (Olympus IX73).

*Measurement of Mitochondrial Membrane Potential (MMP) and ATP level*: Oocytes were incubated with 10 µM Image-iT™ Tetramethylrhodamine (TMRM, Invitrogen, Belgium) in M16 medium at 37°C for 30 min to measure MMP level in oocytes. Subsequently, the stained oocytes were washed and examined using a fluorescence microscope (Olympus IX73). ATP content within individual oocytes was quantified using an Enhanced ATP Assay Kit (Beyotime Institute of Biotechnology, Shanghai, China), following the manufacturer’s protocol. A series of ATP standards were prepared, encompassing a range from 0 to 40 pmol ATP. Oocytes were lysed by treatment with 20 µM lysis buffer in 0.2-ml RNA-free centrifuge tubes, followed by centrifugation at 4°C and 12,000 g for 5 minutes. Unless specified, all procedures were performed on ice. ATP detection solution was dispensed into 96-well plates and incubated at room temperature for 3-5 minutes. In each well, standard solutions and ATP detection diluents were introduced. Samples were subsequently added to the wells, and luminescence was promptly measured using a luminometer (Infinite F200, Tecan, Switzerland). ATP concentrations were determined from the standard curves. The total ATP level was divided by the count of oocytes in each sample to derive the average ATP content per oocyte (pmol).

*Immunofluorescence*: Oocytes/embryos were fixed in 4% (w/v) paraformaldehyde for a minimum of 24 hours at 4°C, followed by permeabilization with 0.5% Triton X-100 in PBS-0.1% PVA at room temperature for 1 hour and washed three times with washing buffer (PBS-0.1% PVA containing 0.1% Triton X-100). Oocytes/embtyos were subsequently incubated in blocking buffer (3% BSA in washing buffer) for 1 hour at room temperature and incubated at 4°C overnight with different primary antibodies (anti-CDX2, 1:500, for trophoblast (TE); anti-Nanog, 1:1000 for inner cell mass (ICM); anti-γH2A.X, 1:100, for DNA double-strand break, abcam, Cambridge, England) diluted in blocking buffer. After thoroughly washed, the oocytes/embryos were further incubated with the appropriate secondary antibody (goat anti-mouse FITC-conjugated antibody, 1:100, goat anti-rabbit Alexa Fluor™ 594-conjugated antibody, 1:100, abcam, UK) for 1 hour at 37°C, then washed four times in washing buffer. DNA was subsequently stained with 4’,6-diamidino-2-phenylindole (DAPI, Vector Laboratories Inc., Burlingame, CA). The stained oocytes/embryos were mounted on glass slides and observed using laser-scanning confocal microscopy (Nikon A1R), with qualitative analysis performed using NIS-Elements AR software (Nikon Instruments).

Oocytes were incubated with 20 mg/ml zeocin in M16 medium at 37°C for 30 minutes to induce DNA damage. Oocytes exhibiting the anti-γH2A.X signal under confocal microscopy were considered as positive cells for DNA damage. The percentage of positive cells and the fluorescence intensity of these positive cells were calculated separately for each group to assess the degree of DNA damage.

*Measuring cytosolic Ca^2+^ and mitochondrial Ca^2+^ alteration:* Cytosolic Ca^2+^ levels were assessed using the cytoplasmic Ca^2+^ probe Fluo-4 AM (Cat. No. 40704ES50, Yeasen, Shanghai, China). The oocytes were processed in M2 medium with 5 μM Flou-4 AM for 20 min and washed three times by DPBS. Subsequently, they were placed within a self-made 1-mm-gap electrode confocal dish (Figure S2 C) and stimulated as previously explained on the stage of a confocal laser scanning microscope (Nikon A1R). Similarly, to assess the dynamics of mitochondrial Ca^2+^ concentration, oocytes were incubated with 5 μM Rhod-2 AM (Cat. No. R1244, Invitrogen) for 30 minutes at 37°C. Fluo-4 AM or Rhod-2 AM were excited using wavelengths of 488 nm or 552 nm and emitted fluorescence was collected at wavelengths of 520 nm or 580 nm.

Images were acquired at 4-second intervals over approximately 30 min and analyzed intracellular fluorescence intensity by ImageJ software. The fluorescence ratio F/F0, defined as the fluorescence intensity (F) of the Ca^2+^ indicator at a specific timepoint during the measurement to normalize to the baseline intensity before exposure (F0), served to illustrate the dynamics of [Ca^2+^]i or intra-mitochondrial Ca^2+^ ([Ca^2+^]mito). In all experiments, nsPEF exposure was initiated following a 1-minute delay to establish a baseline fluorescence level.

*Specific inhibitor treatment*: The ooocytes were incubated in M16 medium with 5 mM EGTA (Yeasen, China) or 10 μM Thapsigargin (TG, MedChemExpress, Shanghai, China) at 37°C for 30 min to deplete Ca^2+^ in the extracellular medium or ER store separately.

Xestospongin C (XC, MedChemExpress, China) is a selective inhibitor of the IP3R on ERs, oocytes were placed in M16 medium with 10 μM XC for 60 min to inhibit IP3R activity. The effect of SOCE in oocytes was inhibited using 2-Aminoethyl diphenylborinate (2-APB, MedChemExpress, China). The oocytes were incubated with 10 μM 2-APB for 30 min in M16 medium.

*In Vitro Fertilization*: Sperm for IVF procedures were obtained from 9-week-old male CD1 mice. The cauda epididymis of the sacrificed was collected and separated with scissors in 100 μL Human Tubal Fluid (HTF) medium. The sperm released from the epididymis were incubated for 1 h at 37°C and 5% CO2 conditions. Cumulus-free oocytes were transferred to 90 µL drops of HTF medium, followed by the addition of 1×10^5^ sperm/ml. After a 15-minute incubation, excess sperm were washed away, and oocytes were either loaded with Fluo-4 AM for [Ca^2+^]_i_ monitoring or cultured in vitro for embryo development as previously described.

*Plasmid Microinjection*: Oocytes were transfected with the GFP-C1-PLCdelta-PH DNA construct, which consists of the Pleckstrin homology (PH) domain of PLCdelta (with tagged IP_3_ or PIP_2_) fused with a green fluorescent protein (GFP) (Addgene plasmid 21179). The microinjection of an optimal plasmid concentration (150 μM) was performed on germinal vesicle (GV) stage oocytes using a microinjection instrument (Eppendorf, Hamburg), followed by a 30-minute incubation period. Subsequently, the oocytes were arrested at the GV stage in M16 medium supplemented with 2.5 μM milrinone for 4 h. Following thorough washing, the oocytes were cultured in M16 medium under mineral oil at 37°C and incubated in a 5% CO2 atmosphere during the GV to MII stages for 12-14 hours. Finally, MII oocytes with GFP-C1-PLCdelta-PH signals were placed in a self-made electrode dish and exposed to 10 ns pulses with medium intensity. The laser scanning fluorescence images were acquired at a rate of 1 image per 4 s for 30min.

*Human oocyte collection*: A total of 9 patients (oocyte donors) underwent controlled ovarian stimulation (COS) using the gonadotrophin-releasing hormone (GnRH) antagonist protocol. The COS was initiated with a dose of gonadotropins (Gonal-F, Merck, Germany; 150-300 IU/day) on day 2-3 of the menstrual cycle. Gn dosages were adjusted according to the ovarian responses evaluated by transvaginal ultrasonography and total serum levels of estradiol (E2) and luteinizing hormone (LH). A GnRH antagonist (Cetrotide, Merck, Germany; 0.25 mg/day) was provided when the leading follicle diameter was 13-14 mm.

Once at least two follicle diameter achieved 18 mm, triptorelin acetate (Decapeptyl, Ferring, Switerland, 0.2 mg) and hCG (Merck, Germany; 6500 IU) was used to trigger follicle maturation. Cumulus-Oocyte Complexes (COCs) were retrieved at 36 h later with ultrasound-guided transvaginal follicular aspiration.

*Human oocyte in vitro maturation*: The immature oocytes of germinal-vesicle (GV) oocytes and meiosis I (MI) stage oocytes were cultured in 20 μl droplets of G-2Plus medium (Vitrolife, Sweden) supplemented with 0.1 IU/ml FSH (Puregon) under paraffin oil. After 24-48 hours cultured at 37℃, 6% CO2, 5% O2, and 89% N2, IVM-MII oocytes were frozen using Cryotop methodology with vitrification kits (Kitazato Corporation, Japan)

*NsPEFs exposure on human oocytes and calcium imaging measurement*: Human thawed oocytes were incubated at 37°C for 30 minutes in G-1Plus medium supplemented with 5 μM Fluo-4 AM. Subsequently, the oocytes were transferred to G-1Plus medium between electrodes and were subjected to nsPEF pulses with different parameters. Due to the different sizes and states of human and mouse oocytes, we stimulated human oocytes with nsPEF at low (20 kV/cm), medium (23 kV/cm) and high (25 kV/cm) intensity with 10-50 pulses, respectively. The intervals between pulses were approximately 1 second.

*Statistical analysis*: All experiments were repeated at least three times. Data are presented as means ± SEM, unless otherwise stated. Statistical comparisons were made with Student’s t tests or one-way ANOVA tests, where appropriate. The *P*< 0.05 was considered statistically significant.

## Supporting Information

Figures S1–S2 and Table S1

Fig. S1 Mitochondria are involved in nsPEFs-induced sustained [Ca^2+^]i oscillations Fig. S2. The utilization of nsPEF generator to oocyte

Table S1. The electrical dose in different dishes by nsPEF stimulation

Video 1 Sustained cytoplasmic Ca^2+^ oscillations induced by nsPEF stimulation at medium intensity, related to Figure 1.

## Acknowledgements

The authors would like to express their gratitude to Dr. Kai Zhang (Beijing Taijie Magnetoelectric Research Institute, Beijing, China), Dr. Jue Zhang (Academy for Advanced Interdisciplinary Studies, Peking University, Beijing, China) and Dr. Jiahui Liu (Academy for Advanced Interdisciplinary Studies, Peking University, Beijing, China) for providing the generator of nsPEF and conversion formula for electrical doses, as well as sharing with us their knowledge and skills regarding the nsPEF device protocol. This study was supported by the National Natural Science Foundation of China (Grant No. 82071715). Yidan Sun and Tong An contributed equally to this work.

## Author contributions

Y.S. participated in study design, execution, analysis and manuscript drafting. T.A. and R.L. participated in patient oocyte collection, in vitro maturation and culture, and revised the manuscript. Y.L., H.X., Y.Z. and X.F. assisted with data analysis and revised the manuscript.

L.F. participated in patient oocyte collection. S.H., Z.S., X.B. and Y.F. assisted patient oocyte collection. Y.-P.H. and Q.L. conceived the idea, planned the study and critically revised the manuscript. All authors read and agreed to the published version of the article.

